# Spatially explicit impact of enhanced leaf litter leaching on the forest soil microbiome

**DOI:** 10.1101/2024.04.11.588997

**Authors:** Simiao Wang, Filippo Miele, Paolo Benettin, Manon Frutschi, Mitra Cattry, Pierre Rossi, Nicolas Jacquemin, Andrea Rinaldo, Rizlan Bernier-Latmani

## Abstract

Climate change is expected to affect precipitation intensity and soil temperature and indirectly impact the release of leached dissolved organic carbon (LDOC) from leaf litter during the early stages of its decomposition, which could affect the health and function of forest soil ecosystem. Here, we experimentally investigate the spatially-explicit impact of LDOC on the forest soil microbiome and the associated biogeochemical processes. Homogenized soil columns were subjected to realistic artificial precipitation for 3 months with the initial level of LDOC adjusted by the number of times the leaf litter was flushed in preparation for the experiment. Hydrological and geochemical parameters (redox potential, pH, dissolved oxygen, soil moisture, matric potential, chemical speciation) were measured continuously as a function of time and depth. The same initial microbial community developed into distinct communities under different LDOC and above and below the water table. The LDOC from leaf litter increased the availability of carbon (C) and nitrogen (N) in porewater four-fold and two-fold respectively in the first two weeks. This resulted in the expansion of the anoxic zone above the water table and a decrease in the soil microbial metabolic potential for cellulolysis and N_2_ fixation in unsaturated soil along with an increase of soil microbial metabolic potential for fermentation at all depths. Finally, increased LDOC decreased the stability, phylogenetic diversity, and complexity of the soil microbiome, limiting its functional diversity. Thus, management of leaf litter should receive more attention due to its indirect role in the impact of climate change on the soil microbiome.

**Highlights:**

- Decreased microbiome diversity and stability due to enhanced leaf litter leaching
- Expansion of the anoxic zone into the unsaturated zone due to increased organic carbon supply
- Decreased soil microbiome metabolic potential for cellulolysis and N_2_ fixation in unsaturated soil
- Depth-dependent response of microbial community to increased organic carbon availability
- Implications for soil response to climate change

## 1 Introduction

Due to the impact of intense human activity starting at the beginning of the industrial period (mid-18th century), a large amount of greenhouse gases has been released to the atmosphere resulting in climate change and a rise in the global surface temperature by 0.95-1.20 °C in 2011-2020 as compared to 1850-1900 (Crutzen, 2006; IPCC, 2021; Trenberth, 2018). In addition to increasing surface temperatures, climate change is also expected to affect global precipitation with an increase in the intensity of precipitation events in western Europe (IPCC, 2021).

Changes in temperature and precipitation could have a series of impacts on forest ecosystems, one of them being their impact on changes in leaf litter leaching that could alter the biogeochemistry of these soils. Although leaching mainly occurs in the early stage of leaf litter decomposition, this process could release 2.5-15% of leaf litter carbon in the form of leached dissolved organic carbon (LDOC), depending on plant type (Hagedorn and Machwitz, 2007; Rizinjirabake et al., 2019; Schreeg et al., 2013). In contrast to the recalcitrant organic carbon fractions, such as lignin, cellulose and hemicellulose that require specialized microbial decomposition, LDOC is readily biodegradable and bioavailable, therefore it may have a non-negligible impact on the forest soil microbiome (Aerts, 1997; Krishna and Mohan, 2017; Olson, 1963). In previous studies, shifts in temperature and precipitation were found to lead to more LDOC in soil from subalpine forests, alpine meadows, and agricultural fields as a result of enhanced root exudation and net primary productivity (Hu et al., 2017; Wu et al., 2011; Yin et al., 2013).

The effect of increased LDOC on the forest soil microbiome could promote a shift in the forest soil microbiome from oligotrophic to copiotrophic taxa, favoring *Acidobacteria, Bacteroidota, Actinobacteria* and *Proteobacteria* (Dou et al., 2023; Fierer et al., 2007; Lladó et al., 2017; Xu et al., 2021), with greater enzymatic activity associated with organic carbon mineralization (Xu et al., 2021). However, such studies usually do not consider the fact that LDOC transported in soil via leaching may impact the soil microbiome in a spatially heterogeneous manner. For example, deep soil with limited organic carbon availability could be more sensitive to LDOC increases as compared to surface soil with high organic carbon availability (Jílková et al., 2020). Investigating the depth-resolved impact of LDOC release on the forest soil microbiome may reveal key controls that are not readily apparent in batch systems.

Here, we conducted a laboratory-scale forest soil column experiment under controlled rainfall conditions (following Poisson stochastic processes) aiming to mimic field conditions as closely as possible. Our goal is to investigate the effect of LDOC released from leaf litter on the spatial distribution, diversity and metabolic potential of the soil microbiome.

We collected and homogenized soil from a forest near Lausanne (Switzerland) and used it to fill three lysimeters 60-cm in height and 30-cm in diameter placed indoors under temperature-controlled conditions. After three months of irrigation reproducing realistic daily precipitation (Rodriguez-Iturbe et al., 1999, 1987), we probed the change in the spatial distribution of an initially uniform microbial community under varying LDOC conditions. The spatial distribution of the microbial community at the beginning and at the end of the experiment informed its compositional trajectory and metabolic potential (Louca et al., 2016) as a function of two environmental factors: available LDOC and depth.

## 2 Material and Methods

### 2.1 Experimental set-up

Experiments were designed to probe the impact of high vs. low leaf litter leaching conditions on a forest soil microbiome using continuously monitored weighted soil columns (also called weighing lysimeters or simply lysimeters, **Figure 1**). Three lysimeters were selected as the experimental set-up because they allow interrogation of the vertical distribution of organic carbon and of the microbiome and provide well-controlled conditions in which only one variable is modified at a time. Classic batch system experiments, in which varying amounts of LDOC are added would not capture vertical gradients and, therefore, would miss what we hypothesized would be a crucial spatial component due to the physical location of the leaf litter above the soil (Fierer et al., 2007; Xu et al., 2021). In contrast to classic batch system experiments, the lysimeters used in this experiment capture spatially explicit changes of biogeochemical characteristics within soil columns.

**Figure 1:**
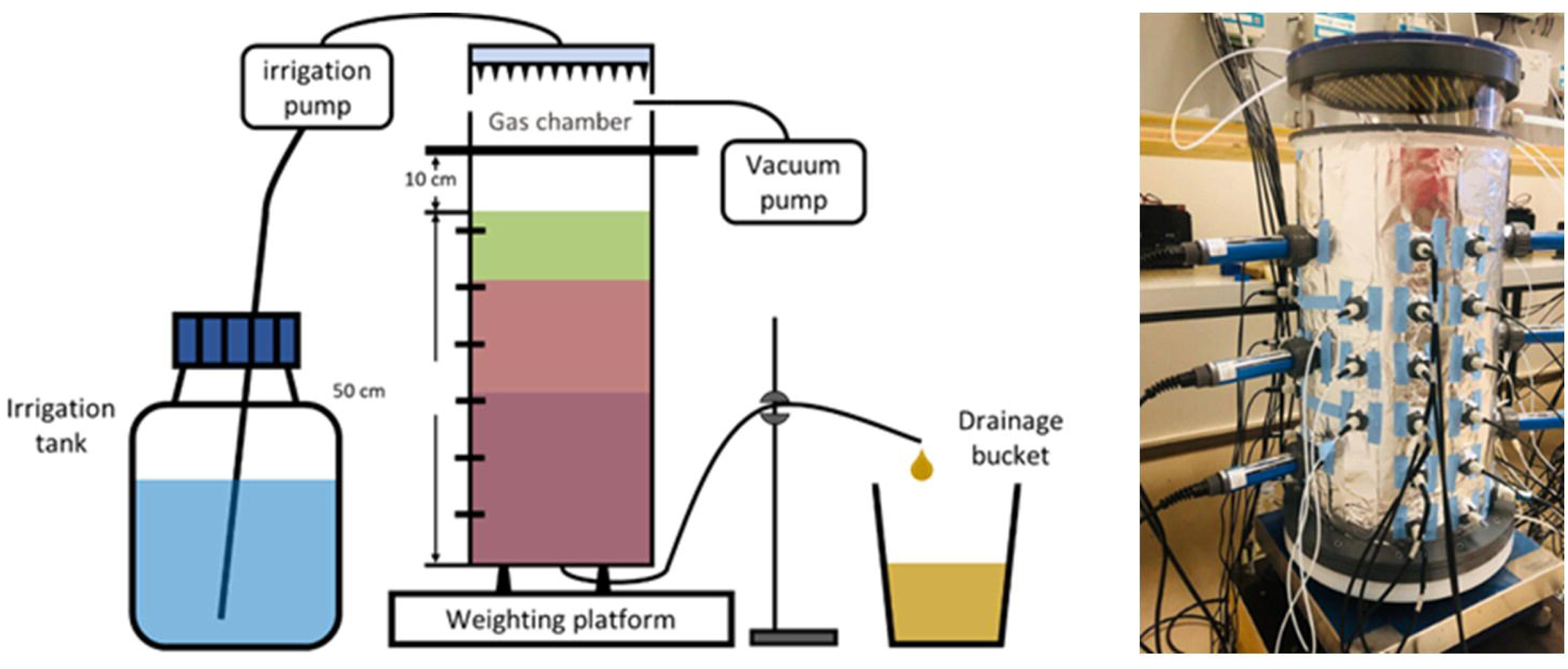
Schematic of the lysimeter set-up (left) and picture of one lysimeter (right). In the schematic plot, the green block represents the soil and leaf litter mixture, and the brown blocks represent forest soil, with the darker brown color to indicate the soil below the average water table.

Our three lysimeters are PVC cylinders (60-cm in height and 30-cm in diameter) whose relatively large size helps reduce wall disturbances. The bottom of each column was equipped with a porous plate, through which porewater drains by gravity. The porewater leaching out of the porous plate was directed through a funnel into a small pipe bending upwards to function as a siphon and to control the water table level. The water table was controlled to remain between 28 and 36-cm below the soil surface. However, at times, due to porous plate clogging, the water table temporarily rose higher.

In September 2021, soil (top 20 cm) was collected from a beech/pine mixed forest near Lausanne (Switzerland), in a site that had been characterized as part of SwissMEX (Mittelbach, 2011). Visible plant material was removed from the soil which was dried at 40 °C for two days, then ground and sieved to 2 mm (Huang et al., 2011). The soil was finally homogenized by mixing all soil batches in a clean cement mixer. Some soil was further sieved to remove particles < 50 µm to produce “coarse soil”. The use of sieved and homogenized soil for the entire lysimeter (rather than trying to capture soil horizons) was a deliberate decision to ensure reproducible hydrological characteristics across replicate lysimeters. Furthermore, it enabled the conclusive probing of the impact of two variables (LDOC and extent of saturation) on the soil microbiome.

Each lysimeter was packed with sieved soil via a dry-packing method. The bottom 5 cm was packed with 4 kg of “coarse soil” to minimize bottom plate clogging (this represents the “initial bottom” soil, 45-50-cm from the surface of the lysimeter). A bentonite clay slurry was painted onto the sides of the lysimeter to minimize preferential flow paths along the lysimeter walls. Subsequently, 4.2 kg of sieved soil was added at a time into the lysimeter and compacted with a wooden compactor to a 5-cm layer. To avoid discontinuity between layers, the soil surface was raked before the addition of a new layer. This process was repeated 7 times until the soil column reached 40 cm in height (this represents the “initial middle” soil, 10-45 cm from the top of the lysimeter).

Leaves collected from the same site were dried (60 degree °C for 48 hours) and contained a total organic carbon (TOC) of about 42.5% by dry mass. They were blended and mixed with sieved soil (native TOC around 2.6% by mass). The mixture, containing 18% TOC by mass, was packed into the top 10 cm of the soil column (2.25 kg of soil-leaf mixture per 5 cm). The mixture served as the organic carbon source for the soil column (this represents the “initial top” soil 0-10 cm from the top of the lysimeter).

Prior to the insertion of probes and suction cups, it is standard procedure to backflush the lysimeter to remove air pockets and to establish a “settled” soil column with limited vertical shear forces (to protect sensor integrity). We leveraged this procedure to investigate the effect of the LDOC level on forest soil microbiome composition. Indeed, lysimeters were pre-treated so that the leaf litter would have either low LDOC (control, n=1) or high LDOC (treatment, n=2) conditions. One lysimeter was flushed with artificial rainwater (recipe in **Table S1**) from the bottom of the lysimeter and subsequently drained from the bottom 7 times to remove a substantial amount of LDOC from leaf litter, resulting in a low remaining LDOC condition (*control*). The other two lysimeters were saturated and drained in the same way but only twice, allowing the leaf litter to retain high LDOC (*treatment*).

Suction cups and various probes were inserted into the lysimeters at various depths (detailed information in **Figure S1**) to allow the collection of porewater samples (suction cups) and the continuous monitoring of soil conditions including redox potential (Eh), pH, soil moisture, water tension (matric potential), and dissolved oxygen (DO). Thus, each lysimeter was divided into five layers of 8 cm and one layer of 10 cm, with each layer being characterized by online data from probes and microbial community analysis at or near its midpoint (**Figure S1b**). The six layers were identical (other than thickness for the bottom one) and named for their midpoint depth: 4cm, 12cm, 20 cm, 28 cm, 36 cm, and 44 cm.

A sprinkler system equipped with 163 irrigation needles was installed at the top of each lysimeter and connected to an irrigation pump to control the timing, amount, and duration of precipitation applied to the lysimeter. Such an advanced irrigation system was necessary to guarantee that precipitation was evenly distributed over the soil surface, with intensities that are not unrealistically high.

Rainfall was applied to the lysimeters according to a distribution that followed a Marked Poisson process for 3 months, which closely reproduces real-world precipitation patterns (Rodriguez-Iturbe et al., 1987) (**Figure S2**). The mean monthly precipitation amount (100 mm) and the mean inter-arrival time (3 days) of the marked Poisson process were selected based on average annual precipitation data for Switzerland from 1991 to 2020 (Federal Office of Meteorology and Climatology MeteoSwiss, 2021).

### 2.2 Sample collection and analysis

During the experiment, porewater samples were collected every 3-4 days for 8 weeks with 10 mL pre-evacuated serum bottles and transferred into an N_2_-filled glovebox (MBraun LABstar Pro, Germany). Inside the glovebox, porewater samples were filtered through 0.22 µm polyethersulfone (PES) filters and analyzed for dissolved organic carbon (DOC) and dissolved nitrogen (DN) with a TOC analyzer (Elementar Vario TOC Cube, Germany).

During the packing stage of the lysimeter, samples of soil and soil-leaf litter mixture were collected to characterize the initial microbiome composition in the top, middle and bottom of the lysimeter respectively. At the end of the experiment, a 3-cm diameter auger was used to collect soil cores along the entire depth of the soil column at 5 positions in each lysimeter (one in the center and four in the periphery) as pseudo-replicates. Each soil core was cut into 4-cm segments, and each segment was transferred to a 50-mL sterile tube (Falcon® 50 mL High Clarity Conical Centrifuge Tubes, USA) and stored at −80 °C. These soil samples were used characterize the soil microbiome composition.

DNA was extracted from soil (200-300 mg per sample) using the DNeasy PowerSoil Pro Kit and a QIAcube Automated Robotic DNA/RNA Purification System (both QIAGEN, USA). The extracted DNA samples were amplified with the PacBio bacterial full-length 16S rRNA gene with barcoded primers in a peqSTAR Thermal Cycler (Peqlab, Germany) and sequenced with the PacBio Sequel II system (Pacific Biosciences, USA). Read Quality was check with FastQC (v0.11.9). Dereplication, error modeling, and ASV inference were performed with DADA2 (v1.30.0). Taxonomic classification was done using the RDP Classifier with the SILVA database (release 138.2). Functional prediction of metabolic pathways was carried out using PICRUSt2 (v2.5.0). The bacterial 16S rRNA gene copies were quantified in triplicate by qPCR performed with a Magnetic Induction Cycler qPCR machine (BioMolecular Systems, Australia) using the primers 338f (5′-ACTCCTACGGGAGGCAGCAG-3′) / 520r (5′-ATTACCGCGGCTGCTGG-3′) with a calibration curve based on known concentrations of *E. coli*.

### 2.3 Quantifying leached DOC from additional backflushing

To quantify the amount of LDOC released from the leaf litter, we performed a separate experiment with dried leaf litter subjected to systematic leaching with milliQ water. Leaf litter (2g) was placed on 0.45µm filters in six replicate filtration units. 60mL of milliQ water was added at a time to fully submerge the leaf litter for 2 minutes. The water was then pumped through the filter into a container, using a vacuum pump. This step was repeated twice for three filtration units and seven times for the other three filtration units. Leachates were collected after each filtration cycle for DOC quantification. The remaining leaf litter on filter was collected at the end of the experiment, freeze dried and quantified for its total organic carbon content.

### 2.4 Statistical analysis

Alpha (α) diversity of microbial communities was calculated using the Shannon Index (Shannon and Weaver, 1949) and the computation was performed with R 4.1.3 using the “Ecological Diversity Indices” function of the vegan package (version 2.6-4). Since this metric is sensitive to large variations in sequencing depth (Schloss, 2024), samples were rarefied to the same number of read counts prior to the analysis using the “Rarefaction Species Richness” function of the vegan package (version 2.6-4). β-diversity between microbial communities was calculated using the Pearson correlation coefficient with R 4.1.3 using the “Correlation, Variance and Covariance” function of the stats package (version 4.1.3) and was further analyzed with the “Hierarchical Clustering” function of the stats package using the “ward.D2” method. Co-occurrence network analysis based on the Spearman correlation coefficient (Spearman correlations with |r| > 0.6 and p<0.05) was used to investigate the co-oscillation of microbial taxa that are present in more than 20% of samples under high and low LDOC conditions (De Vries et al., 2018; Gao et al., 2022; Xiu et al., 2021). The calculated network was imported into Gephi v 0.10.1 for visualization and modularity analysis at 1.0 resolution with edge weight (Bastian et al., 2009; Xiu et al., 2022). The robustness of the co-occurrence network was analyzed with R 4.1.3 using the “Analysis of network robustness” function of the brainGraph package (version 3.1.0)

### 2.5 Accession number

The sequencing data for all samples were uploaded to GenBank database with accession number: PRJNA1246177. This sequence read archive will be released upon publication.

## 3 Results

### 3.1 Impact of LDOC on physicochemical soil characteristics

Repeated leaching of leaf litter resulted in the release of DOC, which we take to represent LDOC here. The overall LDOC in leaf litter was estimated to be 9.35±1.08 mg C/g leaf litter. Two leaching cycles resulted in a cumulative loss of 2.29±0.35 mg C/g leaf litter and corresponded to the high LDOC condition, whereas 7 leaching cycles removed 6.25±0.44 mg C/g leaf litter, representing the low LDOC condition (**Figure 2**). This means that almost three times as much LDOC was lost from leaf litter in the low LDOC control versus the high LDOC condition.

**Figure 2:**
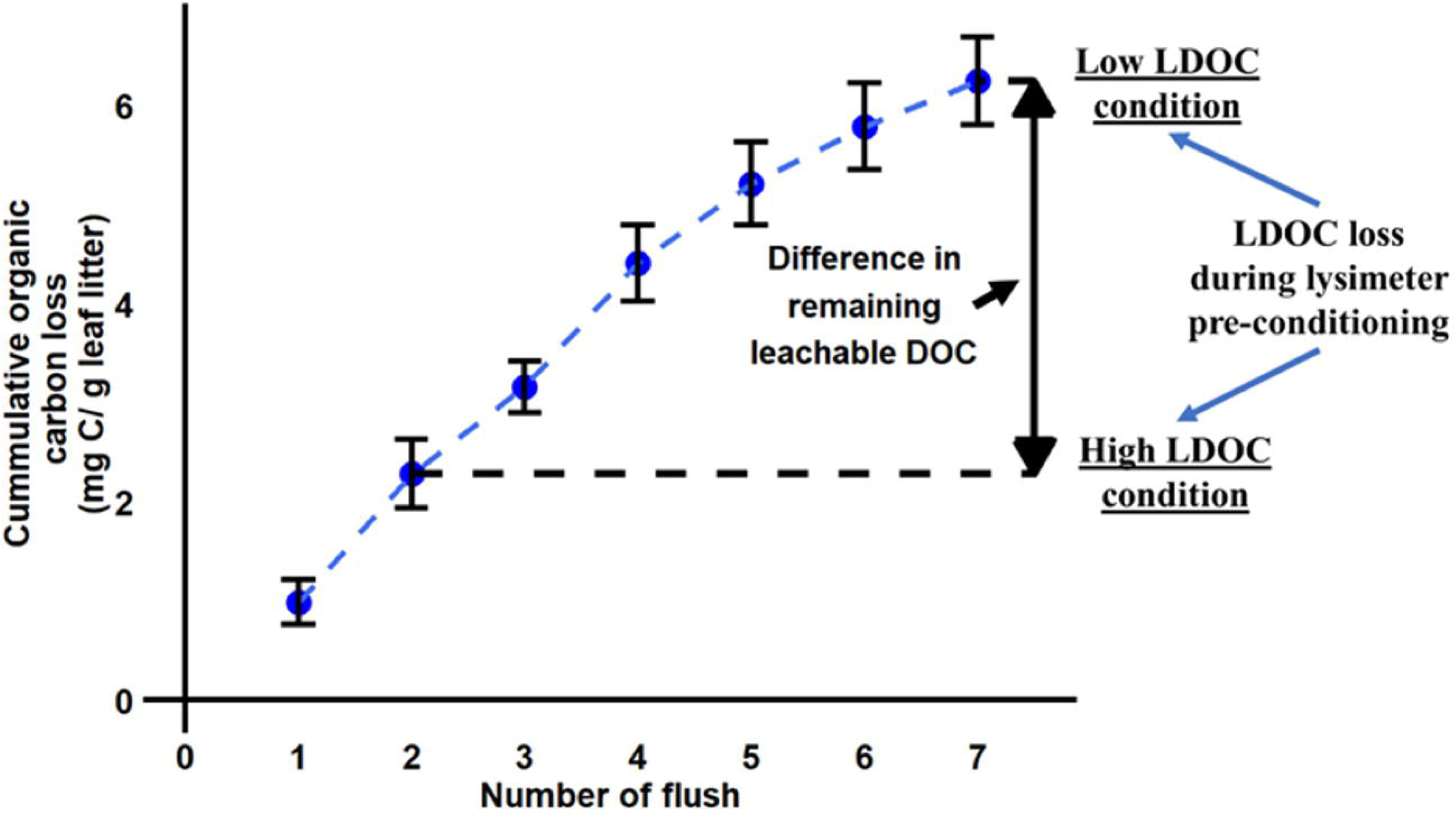
Cumulative organic carbon loss from leaf litter as a function of the number of lysimeter flushes. “High LDOC treatment” and “Low LDOC control” refer to loss in leachable DOC for the two conditions considered here. The High LDOC condition loses less DOC than the low LDOC condition.

The spatiotemporal variation of soil volumetric water content (VWC), Eh, DOC and DN is evidenced in **Figures 3** & **S3**. Fluctuations in soil moisture content at various depths, measured through readings from the volumetric water content (VWC) sensors placed at 4, 20 and 36 cm (**Figure 3d**), were driven by precipitation events and infiltration processes. There were two sharp events with an increase in soil moisture content in high LDOC #2 lysimeter due to clogging events. Except for those, there were no large differences in soil moisture content between treatment and control.

**Figure 3:**
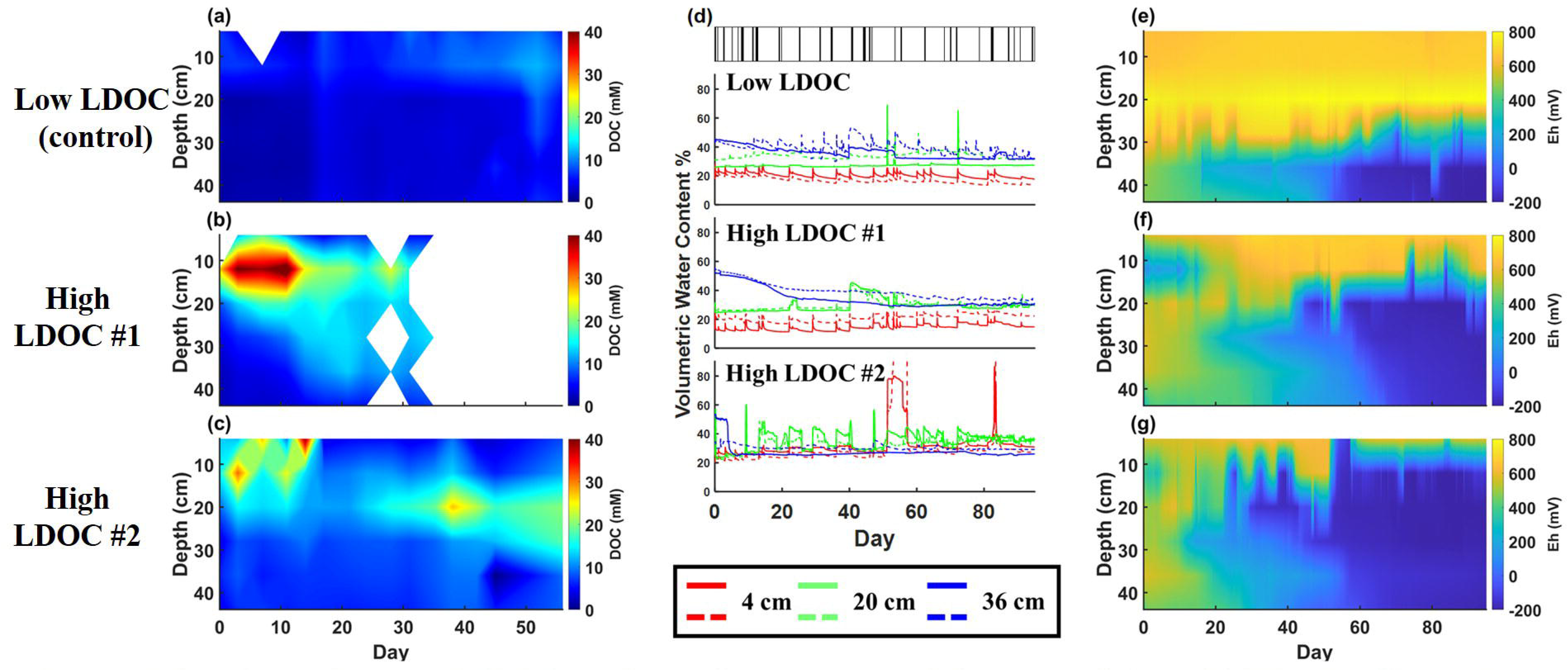
Spatial and temporal response of soil physical and chemical parameters to precipitation events under low and high LDOC conditions. (a)-(c) Porewater DOC concentrations, blank refers to lack of porewater sampling due to low moisture content. (d) Soil moisture content measured as volumetric water content (VWC) at 4, 20 and 36-cm depth. The narrow bar plots at the top represent precipitation events with the bar width proportional to the volume delivered for individual precipitation events. (e)-(g) Soil redox potential values measured at 6 depths every 5 minutes normalized to the Standard Hydrogen reference Electrode (SHE). (a) and (e): Data collected from the lysimeter under low LDOC conditions. (b), (c), (f) and (g): Data collected from lysimeters under high LDOC conditions (High LDOC #1&#2).

Porewater samples were collected from 6 depths from day 0 to day 56 and analyzed for their DOC concentrations (**Figure 3a-3c**). As expected, there was a large difference between the low LDOC control and the high LDOC. In the low LDOC control, only the top two layers (4 and 12 cm) exhibited relatively large DOC concentrations, with the maximum being 12.76±0.20 mM on day 52. This concentration happened simultaneously with a sharp increase in DOC concentration below 12 cm with a peak value reaching 10.34±0.01 mM. In the high LDOC condition, the DOC concentration at 4 and 12 cm reached its highest value (43.74±2.28 mM and 39.52±0.12 mM, respectively) early in the experiment (days 11 and 14) and then stabilized at lower levels (14.97±8.79 mM and 8.42±2.88mM, respectively). Below 12 cm, the DOC concentration increased throughout the experiment. In addition, another DOC concentration peak occurred at 20 cm on day 38, increasing the DOC concentration to 27.32±3.11 mM (**Figure 3c**).

Under low LDOC condition, soil Eh values exhibited *three distinct* responses that varied with depth (**Figure 3e**): (1) At 4, 12 and 20 cm, Eh remained relatively constant between 600 and 800 mV throughout the experiment, reflecting oxidized conditions; (2) At 28 cm, Eh fluctuated between −110 and 700 mV in response to precipitation events, representing the redox fluctuation zone; and (3) at 36 and 44 cm, Eh decreased continuously from 500mV to −200mV with a sharp decrease between days 50 and 60, the reducing zone. In contrast, in the high LDOC condition (**Figure 3f & 3g**), the oxidizing zone was limited to the 4-cm layer that remained at ~600 mV for most of the time, (except for short excursions to −200 mV due to clogging events). In addition, the redox fluctuation zone extended to the 12-cm layer varying between ~100 mV to ~600 mV or −200 and 600 mV (depending on the lysimeter). At day 53, Eh shifted down to between −200 and 200 mV in the high LDOC #2 lysimeter. Finally, the reducing zone extended up to 20 cm for most of the experiment. Therefore, here, the LDOC status has profound implications for the redox state of the soil.

### 3.2 Impact of LDOC on the microbiome

The microbial community composition varied with depth and the LDOC level (**Figure 4**). Five pseudo-replicate cores from each lysimeter showed consistency within treatment and dissimilarity between control and treatment (**Figure S4**).

**Figure 4:**
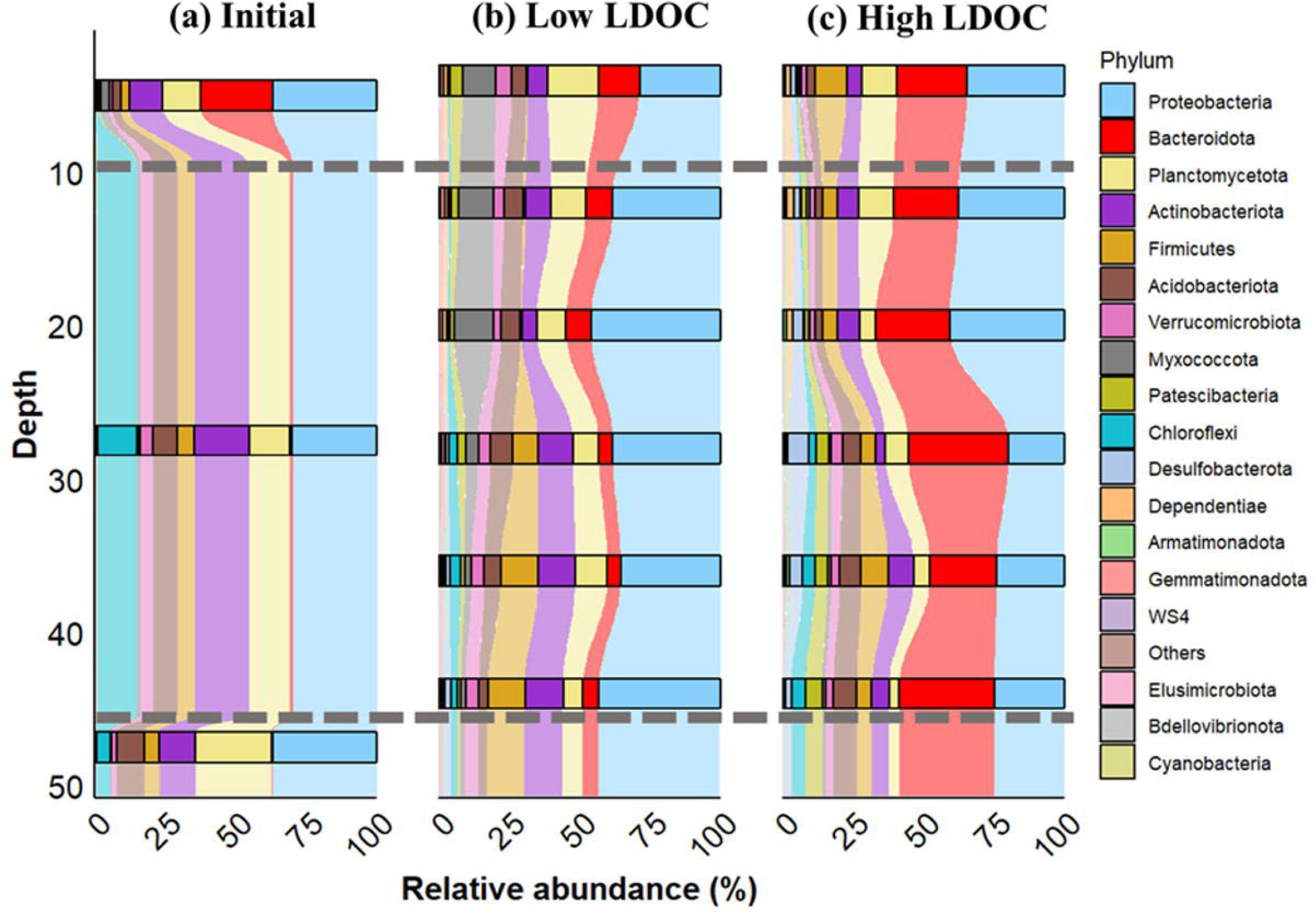
The soil microbial composition at the phylum level based on 16S rRNA gene sequencing as a function of depth. (a) Initial soil microbiome composition during packing; (b) Soil microbiome composition at the end of the experiment under low LDOC condition; (c) Soil microbiome composition at the end of the experiment under high LDOC condition. Phyla with a relative abundance less than 1% were assigned to ‘others’ in this figure. Dashed lines delineate the extents of the three initial packing material across lysimeters.

The initial composition of the soil microbiome (10-45 cm depth) reflects typical forest soil phyla: *Proteobacteria* (29.97 ± 8.02%), *Actinobacteriota* (19.38 ± 0.72%), *Planctomycetota* (14.38 ± 5.40%) and *Chloroflexi* (13.70 ± 6.61%). A similar microbiome was found in the “coarse soil” layer used for the bottom of the lysimeter (**Figure 4a**). The leaf litter and soil mixture (0-10 cm) was distinct and included *Bacteroidota* (25.57 ± 1.85%) and a higher proportion of *Proteobacteria* (36.90 ± 4.53%).

The 3-month experiment resulted in a significant shift in the microbial community under both low and high LDOC conditions (**Figure 4b &c**). For the low LDOC control, the phylum *Myxococcota* appears at 4, 12, and 20 cm (representing 11.61 ± 2.72%, 12.44 ± 4.27% & 13.83 ± 5.54%, respectively), whereas it does not contribute significantly to the microbiome in the high LDOC condition. Additionally, there is a large increase in the contribution of the phylum *Bacteroidota* at all depths in low LDOC (7.32±4.15%) compared to the initial soil (0.60±0.46%) but even a greater increase for high LDOC (27.86±11.74%). Finally, the contribution of the Phylum *Proteobacteria* that is high in the initial soil (33.41±6.72%) remained at the same level in low LDOC at all depths (39.36±9.39%) and high LDOC at 4, 12 and 20 cm (37.60±10.31%) but it significantly decreased in high LDOC at 28, 36 and 44 cm (22.92±6.43%).

The α-diversity of soil microbiomes was calculated using the Shannon Index and plotted as a function of depth (**Figure 5**). Under both low and high LDOC conditions, there is no significant correlation between α-diversity and depth (p = 0.884 for low LDOC condition and p = 0.051 for high LDOC condition). When comparing α-diversity of the same depth from different LDOC conditions, there is only a significant difference in soil microbiomes α-diversity below 12 cm (p <0.01 for 20, 36 and 44 cm and p <0.001 for 28 cm) (**Figure 5**), with a decrease of the α-diversity in the high LDOC case.

**Figure 5:**
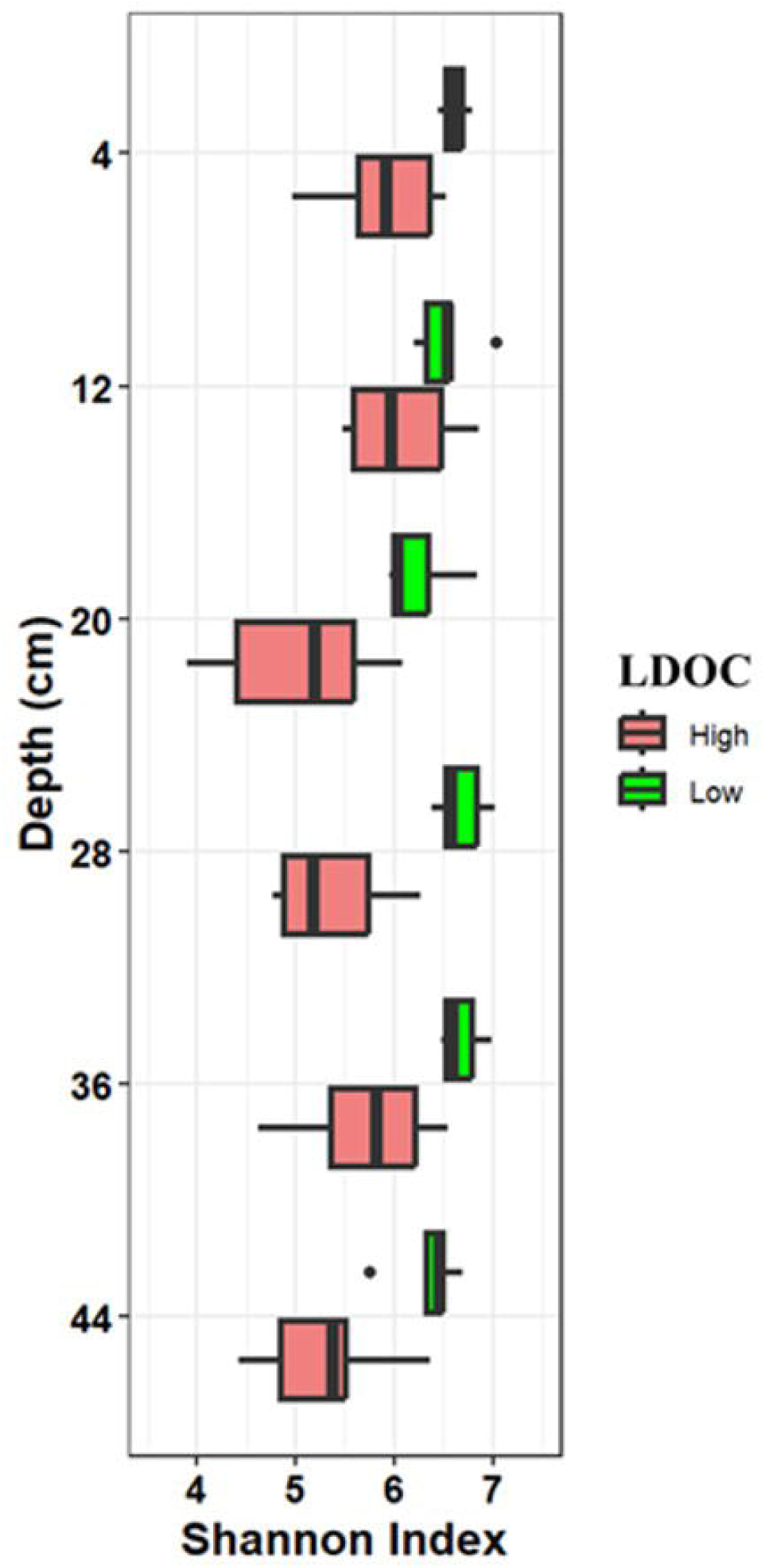
α-diversity (Shannon Index) of rarefied communities as a function of lysimeter depth

Pairwise comparisons of soil microbiomes at different depth and LDOC conditions were performed using the Pearson correlation coefficient for β-diversity (**Figure 6**). When organized according to hierarchical clustering, five sample clusters were obtained based on high within-cluster similarity and lower between-cluster similarity. Except for soil microbiomes collected from the initial samples, soil microbiome compositions clustered according to two factors: depth and LDOC condition. Under the same LDOC condition, soil microbiome composition in samples collected from 4, 12 and 20 cm differed from that of samples collected from 28, 36 and 44 cm. Henceforth, we divide the depth of the lysimeter into two operational parts: shallow (4, 12 and 20 cm) and deep (28, 36 and 44 cm).

**Figure 6:**
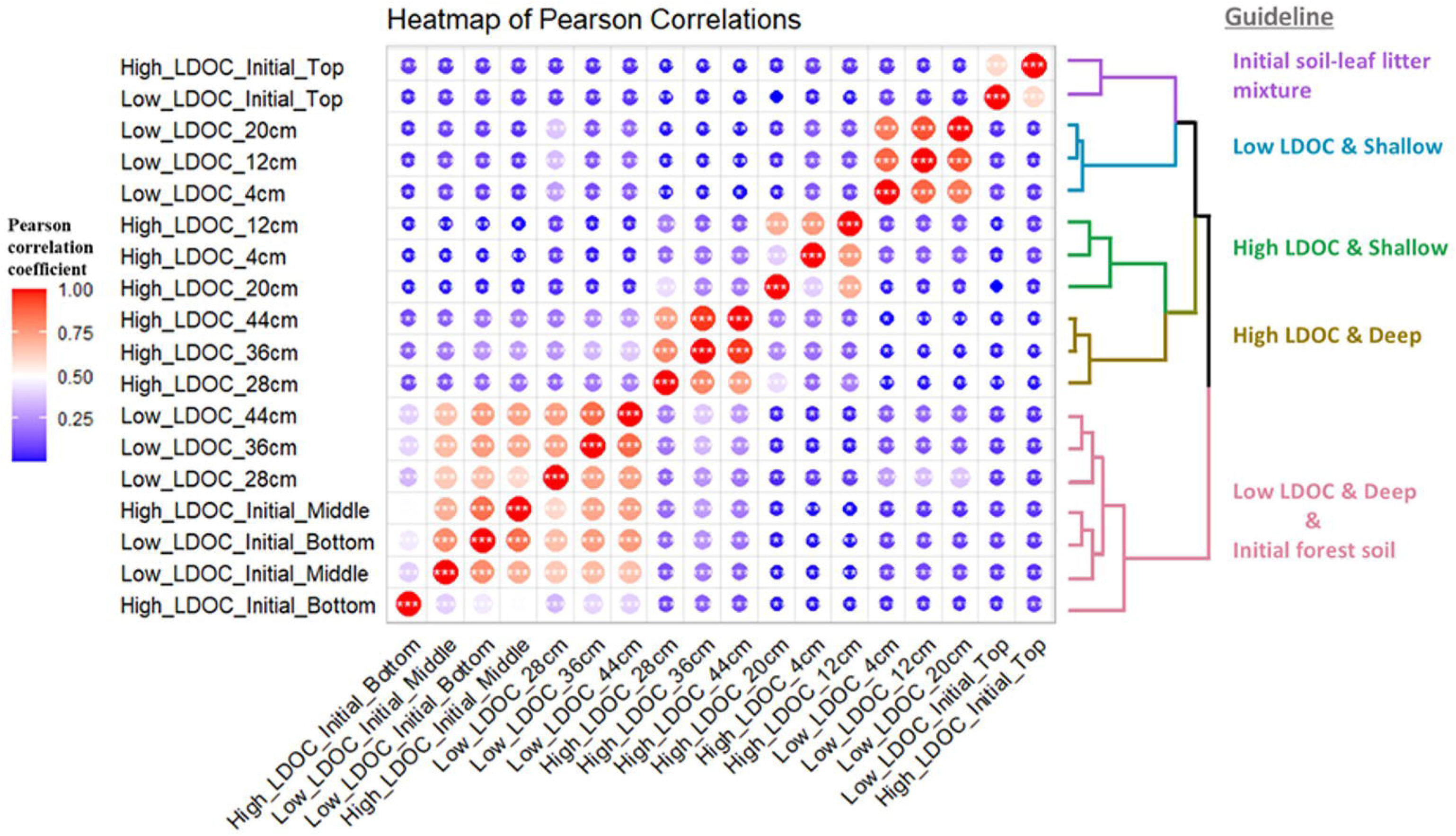
Heatmap of β-diversity (Pearson correlation coefficient) of soil microbial communities. Sample positions are organized according to hierarchical clustering (on the right). Based on the hierarchical clustering, we identified 5 clusters that were labeled based on their location and treatment. The size and color of each circle represent the value of Pearson correlation coefficient comparing the microbiome composition of two samples. Markers in each circle represent level of significance: *: p<0.05; **: p<0.01; ***: p<0.001

To assess the impact of increased LDOC level on soil microbiome composition, a co-occurrence network analysis was conducted at the ASV level for taxa that were present in more than 20% of soil microbiomes collected at varying depths and LDOC conditions (**Figure S5**). This analysis was proven to illustrate the response of the interaction between microbes to various environmental stresses (De Vries et al., 2018; Gao et al., 2022). Key features of the co-occurrence network analysis were extracted and plotted to investigate the effects of the increased LDOC level on the soil microbiome. First, the complexity of the network, which is demonstrated by the number of nodes and edges was evaluated (Gao et al., 2022) (**Figure S6a & S6b**). The number of nodes decreased from 1,022 to 364 and from 835 to 480 when comparing the low LDOC and high LDOC conditions in shallow and deep soil, respectively. A large decrease was also observed in the number of edges: the number of edges in shallow and deep soil decreased from 53,315 and 20,510 under low LDOC condition to 2,278 and 1,466 under high LDOC condition.

Second, the robustness of co-occurrence network was considered. Following the method applied by Liu et al., 50% of the nodes were randomly removed 100 times to calculate the robustness of each co-occurrence network (Liu et al., 2022). Under low LDOC condition, both shallow and deep soil microbiome co-occurrence networks showed strong robustness (both 0.500 ± 0.001). With increased LDOC levels, the robustness of the co-occurrence network decreased to 0.459 ±0.011 and 0.389 ± 0.024 for shallow and deep soil, respectively.

Lastly, the number of keystone taxa was also used to assess the stability of co-occurrence networks (Liu et al., 2022). Keystone taxa are usually defined as highly connected taxa that hold significant influence over community structure and function regardless of their abundance across space and time (Banerjee et al., 2018). In co-occurrence network analysis, they are identified as microbes with high connections (degree > 100) and low betweenness centrality (betweenness < 5,000) (Ma et al., 2016). Under low LDOC condition, 397 and 52 keystone taxa were identified for soil microbiomes in shallow and deep soil (**Figure S6c & S6d**). In shallow soil, most keystone taxa belong to *Proteobacteria* (118 ASVs), *Planctomycetota* (67 ASVs) and *Acidobacteriota* (40 ASVs). While in deep soil, most keystone taxa belong to *Actinobacteriota* (14 ASVs), *Acidobacteriota* (9 ASVs) and *Proteobacteria* (9 ASVs). However, no keystone taxa were identified in soil under high LDOC condition, due to the low number of connections between nodes.

The metabolic potential was predicted for shallow and deep soils under low and high LDOC conditions based on the absolute abundances of the 16S rRNA gene in dry soil via the FAPROTAX database (Gao et al., 2024; Louca et al., 2016) (**Figure 7**). In the shallow soil, an increased LDOC level corresponded to a significant decrease in the abundance of taxa with the metabolic potential for nitrogen fixation, cellulolysis, xylanolysis, photoheterotrophy, and anoxygenic S oxidizing photoautotrophy. On the other hand, it corresponded to a significant increase in the abundance of taxa with a metabolic potential for fermentation, hydrocarbon degradation, NO_3_ ^-^ reduction and ureolysis. In the deep soil, a decrease in the abundance of taxa with a metabolic potential for aerobic chemoheterotrophy, N_2_ fixation, photoheterotrophy, methanotrophy and hydrocarbon degradation was observed for the high LDOC as compared to the low LDOC level. In contrast, the abundance of taxa with a metabolic potential for fermentation significantly increased. In general, increasing LDOC level largely reduced the functional complexity and diversity of the soil microbiome, especially in the deep soil, leading to a change in soil microbiome composition towards a more fermentative microbial community.

## 4 Discussion

The experimental set-up selected for this work has major advantages as compared to batch or field experiments. The controlled gradient approach allows for the systematic investigation of the depth-resolved impact of changes in LDOC on chemical and biological soil properties. Indeed, it enabled the discrimination of soil microbiome shifts as a function of water saturation. Further, the controlled environmental conditions indoors allow for the interrogation of the impact of a single variable on the microbial community. Finally, the extensive instrumentation collecting continuous physicochemical data and discontinuous chemical composition data allows for the detailed monitoring and the observation of spatiotemporal variations in dominant processes.

**Figure 7:**
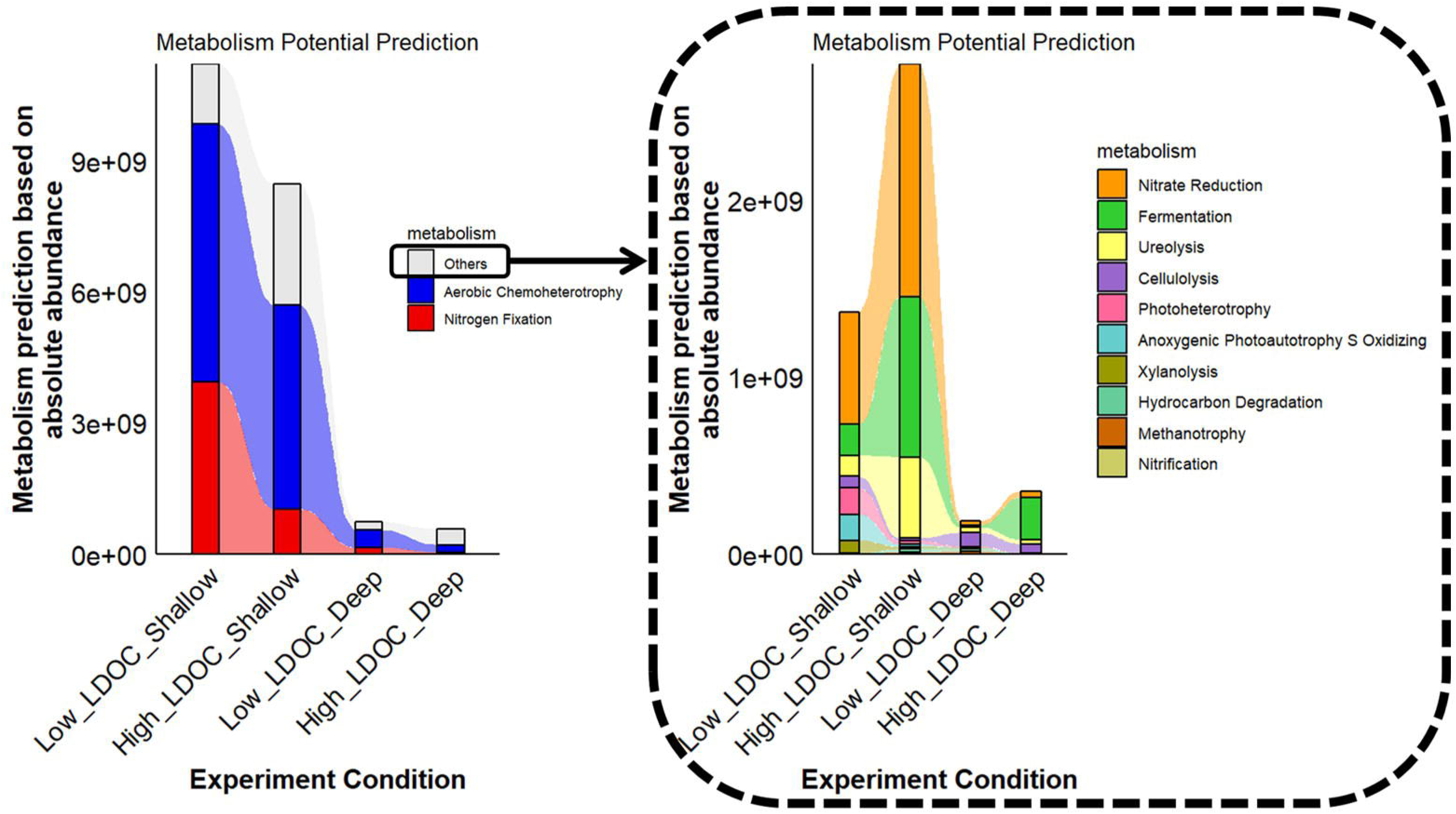
Metabolic potential prediction based on the absolute abundance of 16S rRNA gene copy concentrations in dry soil via FAPROTAX database (Gao et al., 2024; Louca et al., 2016). Metabolism with significant differences between different experimental conditions were selected. Due to the magnitude of differences, aerobic chemoheterotrophy and nitrogen (N_2_) fixation are shown in the left panel while other selected metabolisms are shown in the right panel.

### 4.1 Increased LDOC level results in anoxic zone expansion

The Eh value of 300 mV was used as the operational boundary to distinguish oxic from anoxic environments (Hirano et al., 2013). Under low LDOC conditions, the changes in Eh were consistent with the changes in soil moisture content, with soils at or below 28 cm either reducing or fluctuating in redox (**Figure 3**). This is interpreted as O_2_ entering the soil mainly via hydrodynamic dispersion, which is limited by the degree of saturation (Haberer et al., 2012; Liptzin et al., 2011). With increasing soil water content, O_2_ becomes limited and reductive metabolisms take place. In contrast, under high LDOC, the anoxic zone expanded up to 12 cm below the soil surface, regardless of the water table levels (**Figure 3f & 3g**). In other words, the redox potential decreased to reflect anoxic conditions in the soil despite the lower moisture content. We attribute this to the higher organic carbon availability resulting from LDOC released from leaf litter, especially near the top of the lysimeter (**Figure 3b & 3c**). The large increase in the availability of organic carbon, that here is expected to function as an electron donor, contributes to faster biotic consumption of thermodynamically favorable electron acceptors, such as O_2_ in this unsaturated soil. Upon depletion of O_2_, other redox couples such as Fe(III)/Fe(II) or SO ^2-^/S^2-^set the redox potential, resulting in a decrease in the Eh values. A similar phenomenon was also reported in previous studies in which high organic carbon availability increased the rate of O_2_ consumption and resulted in lower DO and Eh (Husson, 2013). A clear negative correlation between DOC concentration and Eh values can be observed in lysimeters under high LDOC condition (p<0.001, **Figure S7**). Although most of the mass loss from leaf litter leaching typically occurs in the time scale of days or weeks, its impact on the soil redox conditions could be observed for more than 3 months (Jiang et al., 2016). This suggested that the short-term organic carbon release from leaf-litter leaching could elicit long-term effects on soil biogeochemical processes and on the microbiome (see below).

Furthermore, although anoxic conditions were found in soil near the surface and deep within the lysimeter, their impact on soil microbiome composition and metabolic potential differed likely due to the difference in the availability of O_2_ from the atmosphere under these two conditions. Below the water table, O_2_ supply is expected to be limited by diffusion whereas in the organic-rich anoxic zone, O_2_ supply may remain the same but the biotic consumption rate would exceed the supply rate even above the average water table level (Husson, 2013; Miele et al., 2023; Rubol et al., 2013). As shown in **Figure 6**, although soil Eh values at 12 and 20 cm remain indicative of an anoxic zone under high LDOC condition, the corresponding soil microbiome composition clustered with samples collected at 4 cm and for which Eh values corresponded to oxic conditions most of time. This suggested that Eh was not the dominant factor shaping the soil microbiome composition under high LDOC condition. Moreover, as shown in **Figure 7**, the prevalence of the potential for aerobic chemoheterotrophy did not vary significantly for shallow soil under low or high LDOC condition. This further supports the fact that, even under anoxic conditions, the soil microbiome under high LDOC condition could still receive sufficient O_2_ supply from atmosphere at 12 and 20 cm to support a large aerobic chemoheterotrophic community. In essence, the rate of O_2_ supply is inferior to the rate of O_2_ consumption, resulting in negligible O_2_ content while aerobic metabolism is supported. Thus, considering that the experimental design aimed to control the water table between 28 and 36cm from the soil surface, the “shallow” (4, 12, 20cm) and “deep” (28, 36, 44cm) groups obtained by hierarchical clustering in **Figure 6**, provided evidence that this aim was achieved as the “unsaturated” and “saturated” conditions correspond to these two groups. Therefore, we conclude that high LDOC overwhelms O_2_ supply and results in anoxic conditions due to the rapid consumption of O_2_ by the aerobic community stimulated by the organic carbon.

### 4.2 Increased LDOC level results in decreased microbiome diversity

The stability, diversity and complexity of the soil microbiome were analyzed to develop a thorough understanding of the impact of an increased LDOC level. Microbiome *stability* is defined as the *resistance* to community structure change following a perturbation, or as community *resilience*, the ability to rapidly return to baseline following a perturbation-related change (Rykiel, 1985; Shade et al., 2012). Here, stability was measured by the robustness of the co-occurrence network and by the number of keystone taxa as a function of the changing external forcing. Both analyses suggested a large decrease in soil microbiome stability under the high LDOC condition.

Increased LDOC led to a decrease in α-diversity (**Figure 5)** which may be attributed to the fact that extensive readily bioavailable organic carbon inputs from leaf litter leaching would favor copiotrophic taxa that outcompete slow-growing oligotrophic taxa (Fierer et al., 2007). Supporting evidence for this interpretation was found when considering the soil microbiome compositional changes observed in **Figure 4**. According to previous studies, microbial leaf litter decomposition would promote the growth of *Proteobacteria*, *Actinobacteriota* and *Acidobacteriota* in forest soil (Kim et al., 2014; Štursová et al., 2012), which was consistent with the dominant phyla observed in soil samples collected from lysimeters under low LDOC condition (**Figure 4**). However, with increasing LDOC, the soil microbiome was mainly dominated by *Proteobacteria* and *Bacteroidot*a (particularly the latter). As both phyla host copiotrophs that favor nutrient-rich environments, their dominance suggested that they outgrow and outcompete oligotrophic taxa, leading to a decrease in the overall soil bacterial diversity (Fierer et al., 2012, 2007; Huang et al., 2021).

According to the “insurance hypothesis”, microbiome diversity and functional stability are positively correlated, because diversity buffers the impact of environmental perturbations on microbiome function (Yachi and Loreau, 1999). Indeed, a highly functionally diverse microbiome is more likely to include microbes able to survive and recover following an environmental perturbation (Allison, 2004; Downing and Leibold, 2010; Philippot et al., 2021; Yachi and Loreau, 1999). Conversely, a decrease in microbial diversity may result in functional loss following an environmental perturbation.

A decrease in microbial diversity could also lead to a decrease in microbial network complexity, which characterizes the complexity of the relationships between microbes within a microbiome (Coyte et al., 2015; Zhang et al., 2024). This is reflected by the smaller number of nodes and edges in the co-occurrence network under high LDOC as compared to low LDOC conditions (**Figure S6a & S6b**).

The major finding in this section is that high LDOC negatively impacts soil microbial diversity and microbial network complexity, as shown through several lines of evidence, which suggested that under the projected climate change scenarios, the LDOC input from leaf litter leaching should be controlled to avoid soil microbiomes become more vulnerable to environmental disturbance.

### 4.3 Increased LDOC level results in microbiome metabolic shift

As the complexity of the soil microbial community has been shown to be correlated to its function, decreased complexity could lead to decreased microbial function within the soil and decreased resistance to environmental change (Chen et al., 2022; Luo et al., 2023; Zhang et al., 2024). A recent study has reported that a decrease in soil functional diversity was observed in an increasingly less complex microbial community with increased aridity (Zhang et al., 2024); Furthermore, less complex fungal communities, induced by grass degradation, have been found to be associated with a decreased ecosystem functional diversity (Luo et al., 2023; Zhang et al., 2024). Such changes would be detrimental to ecological function. In addition, the functional complexity of the soil microbiome was proven to play an important role in the persistence of organic carbon, further highlighting its ecological importance in reducing carbon loss from soil (Lehmann et al., 2020). Indeed, we observe that increased organic carbon availability has a profound impact on microbial functional diversity in the saturated soil (**Figure 7)** through the decrease in abundance of seven metabolic pathways and the increase only in fermentation.

In addition to the effect on functional diversity, the increased LDOC level also resulted in significant changes in metabolic potential in both unsaturated and saturated soil (**Figure 7**). Most of the significantly altered metabolic pathways are related to either C or N cycling. We attribute those changes to the large amount of nutrients released from leaf litter via leaching as LDOC.

#### C cycling

Since there is no exogenous organic C added to lysimeters throughout the experiment, the only source of DOC is leaf litter via either microbial decomposition or leaching. As cellulose is one of the most abundant compounds in leaf litter and is less recalcitrant than lignin, the cellulolytic metabolic potential could represent the bacterial leaf litter degradation capability within the lysimeter (Khan and Ahring, 2019; Štursová et al., 2012). Under high LDOC condition, a significant decrease in cellulolytic potential was observed in the unsaturated soil, supporting the suggestion that the soil microbiome is less dependent on bacterial leaf litter decomposition to obtain organic carbon than in the low LDOC condition. As LDOC released from leaf litter was likely readily bioavailable dissolved organic carbon, it would be used preferentially relative to more complex leaf litter carbon as an energy and carbon source. Moreover, the contribution of fermentation to metabolic potential significantly increased at all depths under high LDOC condition, which could be attributed to a combination of the high organic carbon available to microbes and the anoxic environment resulting from the associated high O_2_ consumption rate.

#### N cycling

Due to the N-deficiency in lignocellulosic biomass in leaf litter and the low amount of N applied to the lysimeter via irrigation, bacterial N_2_ fixation, which utilizes atmospheric N_2_ to produce bioavailable N, is expected to play an important role in the bacterial decomposition of lignocellulosic biomass (Harindintwali et al., 2022). This explains the high contribution of N_2_ fixation to the metabolic potential in unsaturated soil under low LDOC condition. However, with increasing LDOC, the metabolic potential of N_2_ fixation significantly decreased in unsaturated soil. This occurred simultaneously with a significant increase in the metabolic potential of other N-consuming biogeochemical processes (NO_3_^-^reduction and ureolysis) in the same zone. These changes suggest that potentially bioavailable N is released simultaneously with LDOC. Berg and Staaf have reported that leaf litter was capable of accumulating N through its binding to lignin and humification products (Berg and Staaf, 1981). N could be released starting from the beginning of leaf litter decomposition via leaching. Thus, the release of LDOC also increased bioavailable N previously bound to leaf litter. Such an explanation is consistent with the high DN concentration observed at the beginning of the experiment (**Figure S3**).

Overall, there is a clear impact of high LDOC on the metabolic potential of the microbiome, with a shift from cellulolysis to fermentation and from N_2_ fixation to nitrate reduction and ureolysis. Increased precipitation intensity expected in Europe based on global climate models (IPCC, 2021) will result in high fluxes of C and N from leaf litter to soil over short periods of time. Based on this study, the shift from low to high LDOC has a lasting impact on the microbial community and its metabolic potential. A shift away from cellulolysis and N_2_ fixation is deleterious to the ability of the soil microbiome to obtain C from recalcitrant leaf carbon and from atmospheric nitrogen. Thus, overall, this change would imply slower terrestrial C cycling and reduced N_2_ fixation. Considering the decreased capability of forest soil microbiome to obtain C and N from environment, the overall productivity of forest soil may be expected to decrease.

In addition, as shown in this study, the response of soil microbiome to enhanced LDOC level varies along soil depth or saturation level, which suggests the necessity of depth-resolved analysis for soil studies. Thus, for future studies of the impact of climate change on soil, it is recommended to collect soil samples at various depths and/or saturation conditions to better constrain the spatial heterogeneity of the impact

## 5 Conclusion

An elaborate lysimeter experiment was conducted to investigate variable leaf litter leaching as a potential control on biogeochemical soil processes due to the expected impact of climate change on LDOC fluxes. The short-term release of LDOC leached from leaf litter increased the availability of C and N in porewater, which enhanced the microbial O_2_ consumption rate and led to a persistent expansion of the spatial extent of the anoxic zone (for about two months). In terms of the soil microbiome, increased LDOC led to a distinct shift in unsaturated and saturated soil via the promotion of *Bacteroidota* growth at all depths and the decrease in the contribution of *Myxococcota* in unsaturated soil and *Proteobacteria* in saturated soil. Overall, it significantly decreased microbial diversity, complexity, and stability by promoting the growth of copiotrophic taxa and by outcompeting slow-growing oligotrophic taxa. This would result in a soil microbiome that is more vulnerable to environmental disturbance and less efficient at retaining organic carbon in soil. Moreover, changes in the soil microbiome also affected its associated metabolic potential. Increased C and N availability reduced the dependency on cellulolysis and N_2_ fixation in unsaturated soil, reducing the capacity of soil microbiome to obtain recalcitrant C and atmospheric N from environment. The metabolic potential of fermentation also increased at all depths in response to expanded anoxic zones and high organic carbon availability. As a result of decreased complexity of the soil microbiome, its functional diversity also decreased, especially for soil below the stable water table level.

This work highlights the fact that short-term enhanced leaf litter leaching could cause a long-term effect on biogeochemical processes in forest soil. Moreover, this would lead to increased C and N loss from forest ecosystem and potential decreases of soil productivity. And thus, leaf litter management strategies should be considered including solutions such as regular collection of leaf litter to control its amount, especially in rainy season. In addition, depth-resolved data enabled the observation of the spatially explicit impact of increased LDOC on forest soil according to the saturation level. This emphasized the spatial heterogeneity of soil microbiome’s response to environmental changes and the need for depth-resolved analysis for future studies of the impact of climate change on soil.

This work points to the indirect impact of climate change on the forest soil microbiome and the potential for its decreased resilience. More investigations are needed to evaluate whether plants would exacerbate or diminish the impact of increased organic carbon input into the soil microbiome and to directly probe microbiome stability under higher LDOC fluxes.

## Acknowledgements

The authors acknowledge key funding provided by the Swiss National Science Foundation through its Sinergia grant CRSII5_186422 and NCCR Microbiomes Grant number 225148. Many thanks to Colin Volet, Jantina van der Meer and Natalia Andrea Montoya Arévalo for their help with the experimental process, Samir Simon Suweis and Wei Xiu for their valuable suggestions on the microbial analysis, and Jana von Freyberg for providing the rainfall composition data. Lastly, we would like to extend our sincere thanks to the Central Environmental Laboratory and the Mass Spectrometry – Elemental Analysis platform at EPFL and the Lausanne Genomic Technologies Facility at the University of Lausanne for sample analysis and sequencing.

## Data availability

All scripts and Dockerfiles used for the sequencing analysis are available at: https://github.com/nlmjacquemin/emlexp104 Geochemical data for all figures were uploaded to Zenodo database (Digital Object Identifier: 10.5281/zenodo.15209708) (Wang et al., 2025b)

