## Supplementary material for "Spatially explicit impact of enhanced leaf litter leaching on the forest soil microbiome": SI Table

| **Compound** | **Concentration (µM)** |
| --- | --- |
| Ca^2+^ | 54.49 |
| Mg^2+^ | 4.79 |
| Na^+^ | 7.24 |
| K^+^ | 2.37 |
| NO_3_^-^ | 7.24 |
| SO_4_^2-^ | 4.79 |
| Cl^-^ | 2.37 |
| HCO_3_^-^ | 108.98 |

Table S1: Artificial rainwater recipe adapted from an unpublished investigation by Jana von Freyberg
