## Supplementary material for "Spatially explicit impact of enhanced leaf litter leaching on the forest soil microbiome": SI Figures

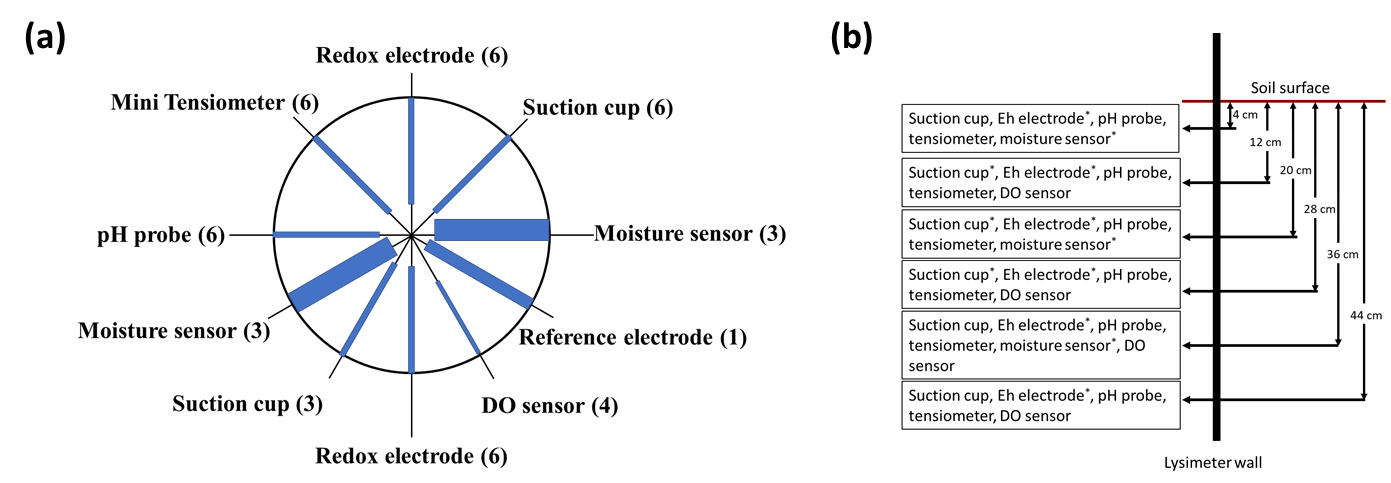

Figure S1: Illustration of the experimental setup. (a) Top view of probes and suction cups within the lysimeter. The numbers in parentheses refer to the number of probes/suction cups in a vertical column; (b) vertical sketch of probes and suction cups within the lysimeter, * refers to duplicate probes/suction cups at the same depth.

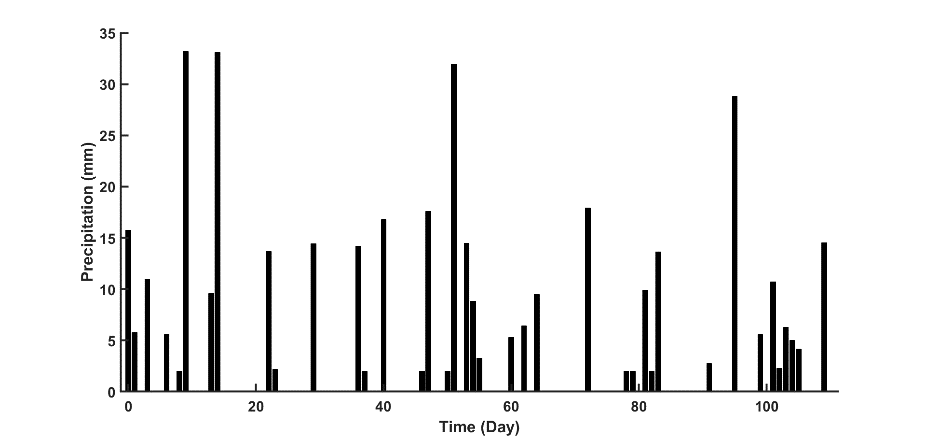

Figure S2: Experimental daily rainfall timeseries generated following a Marked Poisson process (Rodriguez-Iturbe et al., 1987) based on precipitation statistics from Switzerland from 1991 to 2020

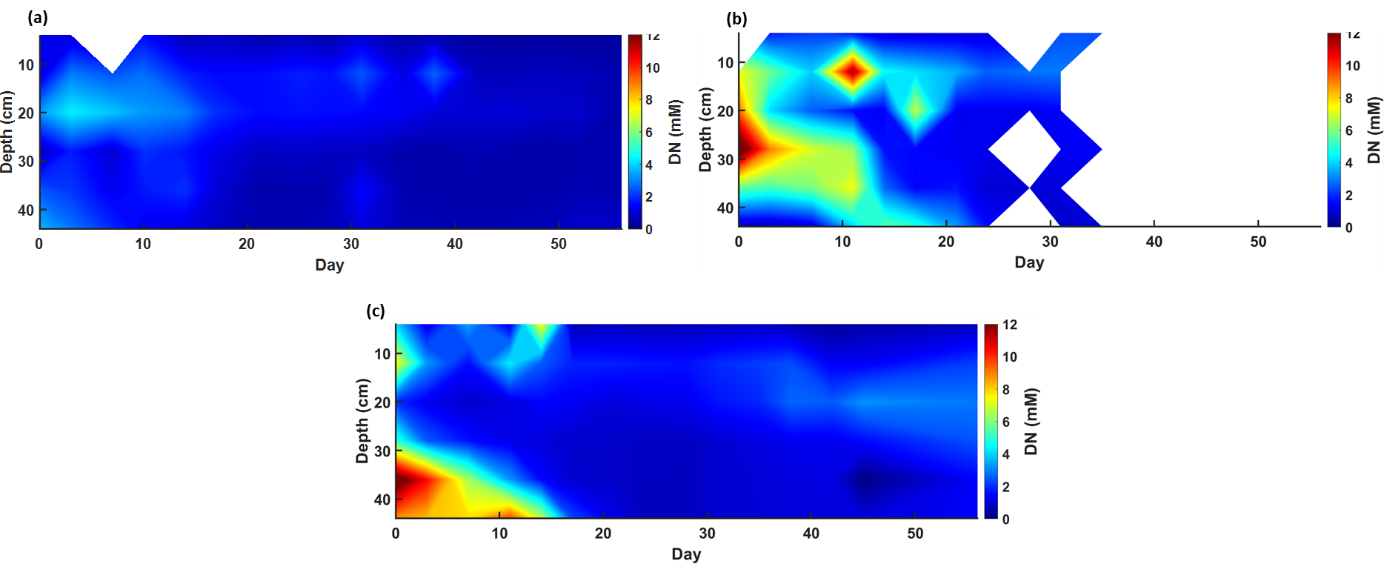

Figure S3: Porewater dissolved nitrogen (DN) concentrations, blank refers to failure of porewater sample collection. (a) refers to data collected from lysimeter under low LDOC condition; (b)&(c) refer to data collected from lysimeters under high LDOC condition

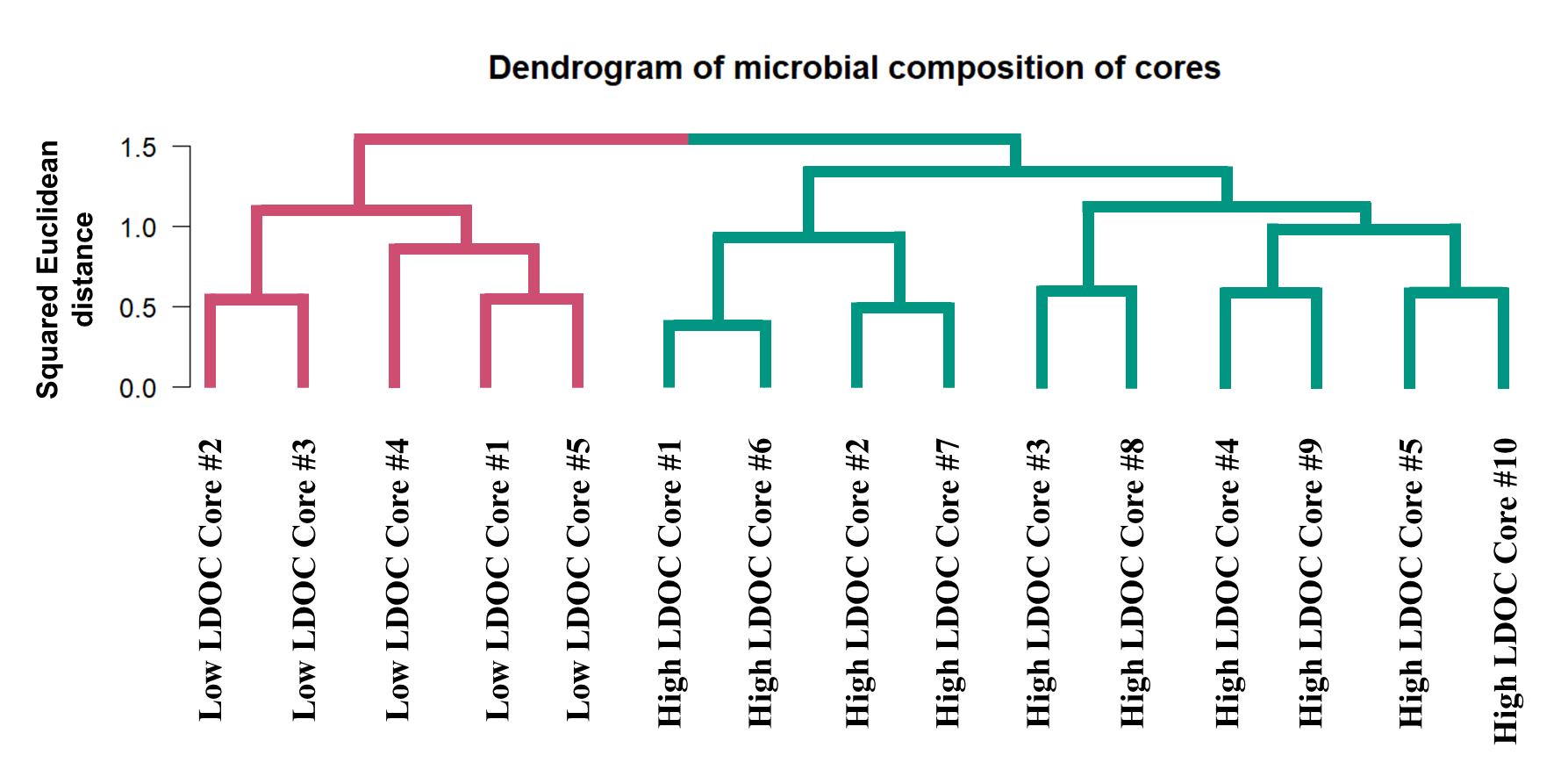

Figure S4: Dendrogram of microbial composition of each core collected from lysimeters. Different colors indicate clustering of samples. The microbial composition of samples showed high similarity within clusters and low similarity between clusters.

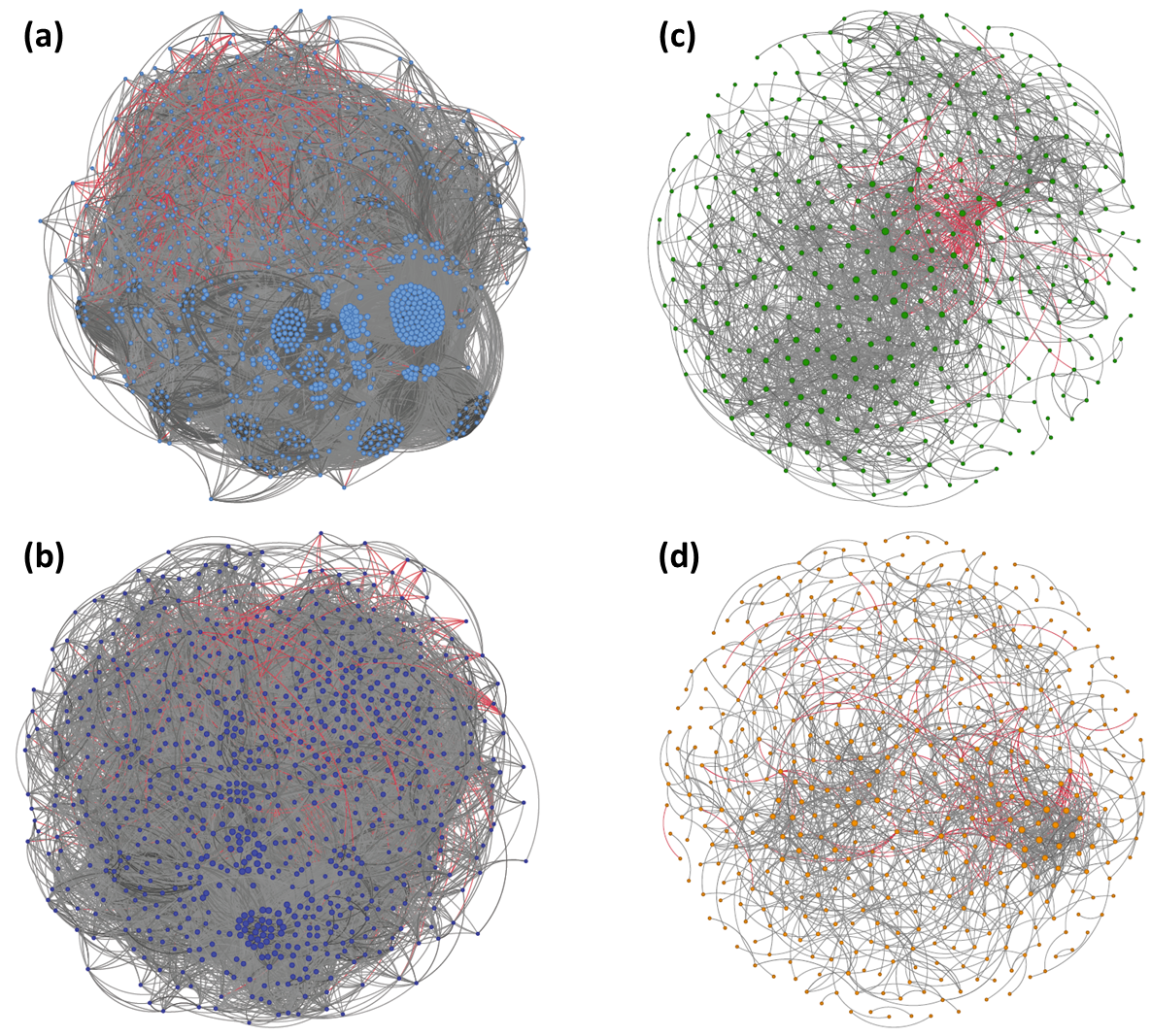

Figure S5: Co-occurrence network analysis of genera present in over 20% of samples (Spearman correlations with |r| > 0.6 and p<0.05). Each node represents one genus, and the size of the node is proportional to the number of connections to this node. Grey connections indicate positive correlations and red connections indicate negative correlations. (a) Shallow soil under low LDOC condition; (b) Deep soil under low LDOC condition; (c) Shallow soil under high LDOC condition; (d) Deep soil under high LDOC condition

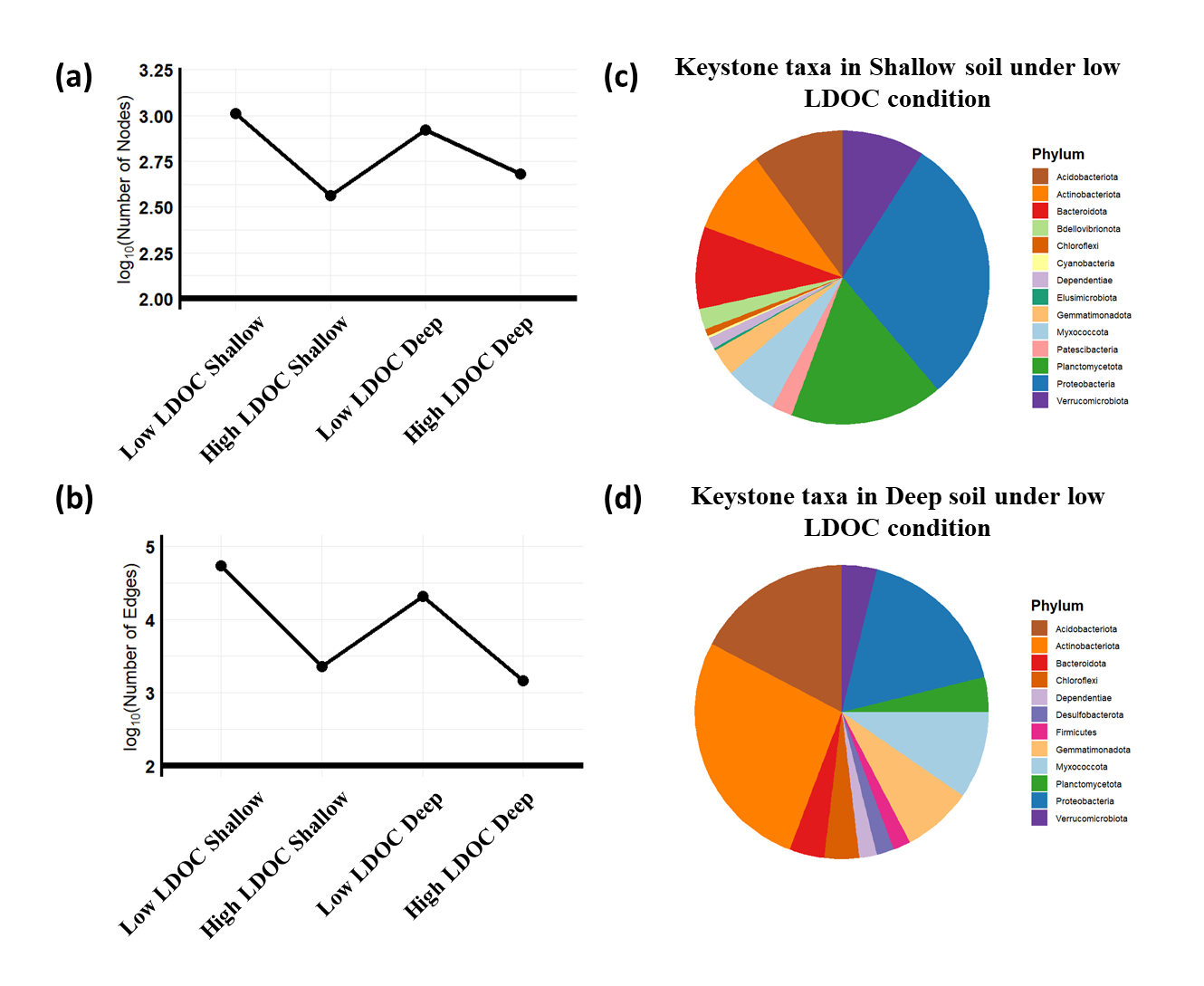

Figure S6: Characteristics of co-occurrence network analysis (Figure S5). (a) number of nodes in co-occurrence networks under different environmental conditions; (b) number of edges in co-occurrence networks under different environmental conditions; (c) keystone taxa composition at phylum level for co-occurrence network of Shallow soil under low LDOC condition; (d) keystone taxa composition at phylum level for co-occurrence network of Deep soil under low LDOC condition

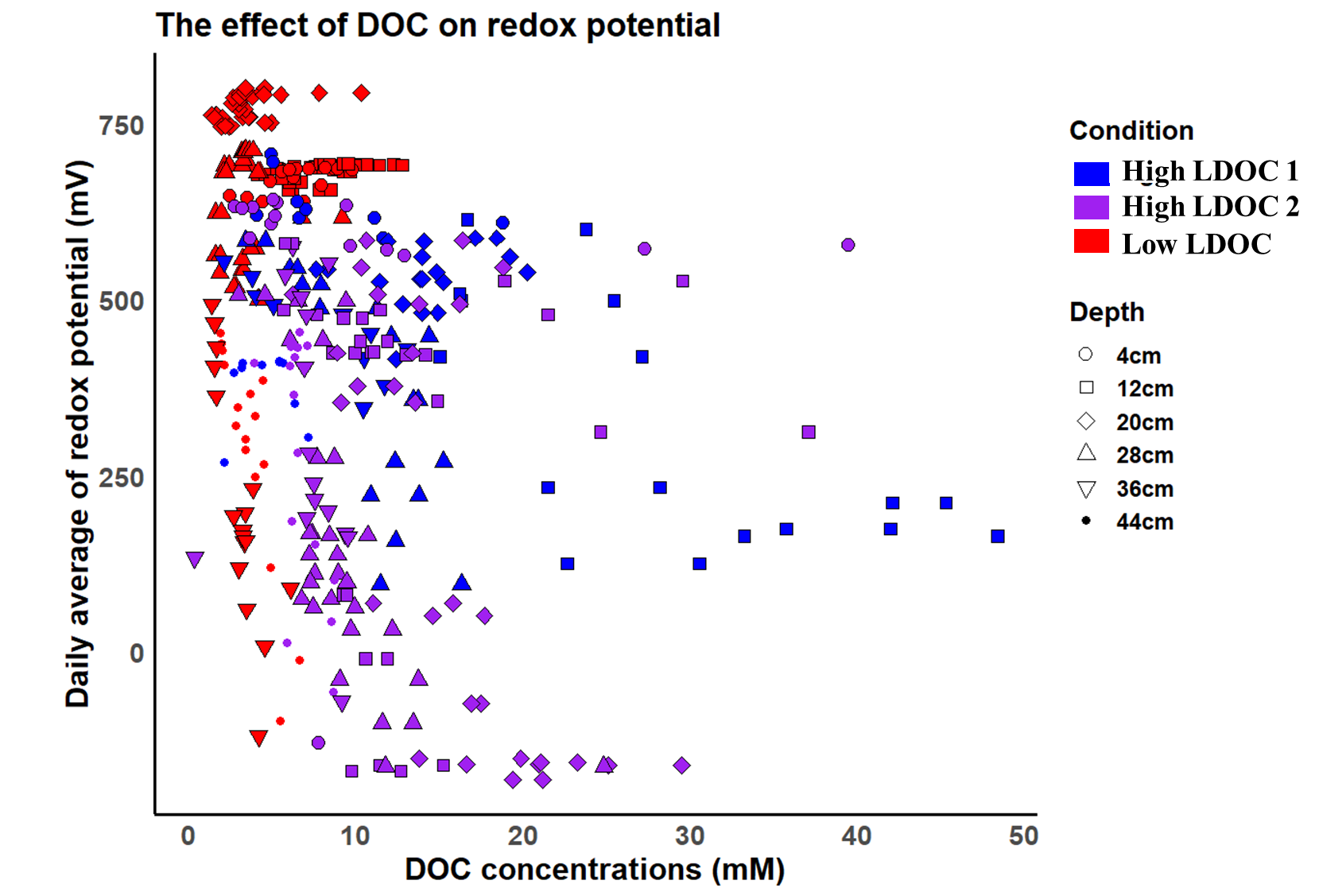

Figure S7: The relationship between DOC concentrations and the daily average redox potential value on the day of sampling.
